# Nucleobase clustering contributes to the formation and hollowing of repeat-expansion RNA condensate

**DOI:** 10.1101/2021.11.08.467691

**Authors:** Ying-Xue Ma, Hao-Zheng Li, Zhou Gong, Shuai Yang, Ping Wang, Chun Tang

**Affiliations:** Innovation Academy for Precision Measurement Science and Technology, Chinese Academy of Sciences, Wuhan, Hubei 430071, China; Beijing National Laboratory for Molecular Sciences, College of Chemistry and Molecular Engineering, and Peking-Tsinghua Center for Life Sciences, Peking University, Beijing 100871, China; Email: (C.T.); Britton Chance Center for Biomedical Photonics, Wuhan National Laboratory for Optoelectronics-Huazhong University of Science and Technology, Wuhan, Hubei 430074, China; E-mail: (P.W.)

**Keywords:** phase separation, Raman spectroscopy, RNA, base stacking, repeat expansion

## Abstract

RNA molecules with repeat expansion sequences can phase separate into gel-like condensate, and this process may lead to neurodegenerative diseases. Here we report that in the presence of Mg^2+^ ion, RNA molecules containing 20×CAG repeats coacervate into filled droplets or hollowed condensate. Using hyperspectral stimulated Raman spectroscopy, we show that RNA coacervation is accompanied by the stacking and clustering of nucleobases, while forfeiting the canonical base-paired structure. At an increasing RNA/Mg^2+^ ratio, the RNA droplets first expand in sizes, and then shrink and adopt hollow vesicle-like structures. Significantly, for both large and vesicle-like droplets, the nucleobase-clustered structure is more prominent at the rim than at the center, accounting for the rigidification of RNA droplets. Thus, our finding has broad implications for the general aging processes of RNA-containing membrane-less organelles.

---

Liquid-liquid phase separation (LLPS) has been increasingly recognized for its relevance in a multitude of biological processes,^[1]^ which enables molecules to form membrane-less organelles. Aberrant phase separation, on the other hand, would lead to the formation of pathological amyloids, hydrogels, or amorphous aggregates, often implicated in neurodegenerative diseases.^[2]^ The research on LLPS has been largely focused on proteins comprising intrinsically disordered or low-complexity regions.^[2a,3]^ Yet, LLPS also occurs between proteins and RNAs.^[1f,4]^ Moreover, repeat-expansion RNAs alone may undergo phase separation without the involvement of proteins.^[5]^ Significantly, RNA condensates usually age rapidly and form gel-like structures with little mobility and fluidity. Therefore, the phase separation of RNA molecules, as opposed to the RNA-encoded proteins, has been proposed to be the etiology of Huntington’s, amyotrophic lateral sclerosis, and other neurodegenerative diseases.^[5]^ Intuitively, the formation of RNA condensate should involve base-paired structure between the repeat-expansion sequences.^[5a]^ Yet, little structural information is available for the RNA assembly in the condensed phase.^[6]^

Various spectroscopic methods have been utilized to characterize the structures of the macromolecules partitioned in the condensed phase. They include fluorescence correlation spectroscopy (FCS),^[7]^ single-molecule Förster resonance energy transfer (FRET),^[8]^ nuclear magnetic resonance (NMR),^[9]^ and Raman scattering spectroscopy^[9b,10]^. The use of fluorescence methods first requires covalent attachments of fluorophores, which may perturb the conformation of the subject molecule. In addition, the photophysical properties of the fluorophore may differ between dilute and condensed phases,^[11]^ complicating the estimation of condensed-phase properties. NMR spectroscopy in principle can provide a detailed structural picture of the macromolecules in the condensed phase.^[9a]^ Yet successful acquisition of NMR data requires the coacervated macromolecules remain fluid and flexible in the droplets, which is often not the case.

Spontaneous Raman scattering spectroscopy arises from intrinsic molecular vibrations. As a result, the spectral features of Raman are not affected by the aggregation state of the macromolecule.^[12]^ Moreover, Raman characterization is label-free and therefore is readily applicable for the coacervated macromolecules. Indeed, the method has been used to evaluate the concentration of ataxin-3 protein^[10]^, and to demonstrate the lack of any structural change of FUS protein in the condensed phase.^[9b]^ However, spontaneous Raman spectroscopy cannot provide temporal information about macromolecular structural changes nor has sufficient spatial resolution for spectral imaging of the macromolecules in a droplet.

On the other hand, hyperspectral stimulated Raman microscope (SRM)^[13]^ allows us to explore structural and molecular basis for the formation of RNA condensate (See methods and apparatus of SRM in Supplementary information, **Figure S1**). As a label-free and molecular imaging methods, SRM have been employed in a wide range of applications in biomedical imaging field.^[14]^ By fingerprinting the molecular vibrations, SRM is capable of identifying and mapping specific covalent bonds with sub-μm resolution,^[15]^ thus well-suited for non-invasive and selective quantification of RNA structure in the droplets.

Previously, LLPS has been observed for RNA molecules with >31×CAG repeats.^[5a]^ We found that an RNA molecule with 20×CAG repeats can already phase separates, which occurs in the presence of a large excess of Mg^2+^ and is consistent with a recent report.^[16]^ Phase separation can be observed for the RNA molecule at as low as 9 μM in concentration, while an increased RNA concentration promotes LLPS in both dimension and the number of the RNA droplets (**Figure 1a**).

**Figure 1.**
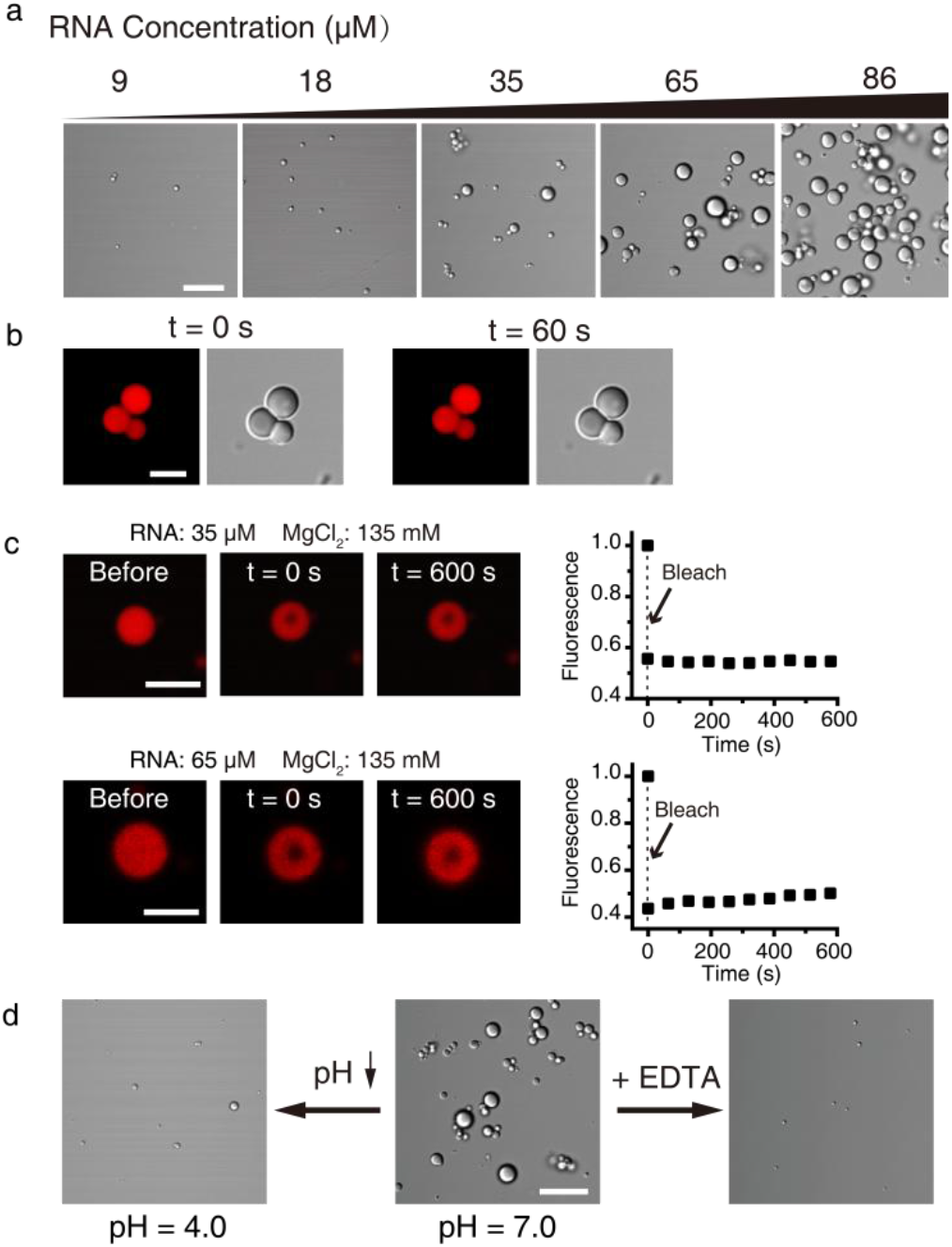
The 20×CAG repeat RNA oligonucleotide phase separates and forms gel-like condensate. **a** DIC images for the RNA droplets formed an increasing RNA concentration. The buffer contains 10 mM Tris•HCl at pH 7 and 135 mM MgCl_2_. **b** The RNA droplets failed to fuse during the time of observation. **c** Fluorescence recovery after photobleaching (FRAP) measurement of the RNA droplets. Photo-bleaching of a droplet in the center, shows a lack of fluorescence recovery. Upper panel: FRAP measurement of the droplet formed by 35 μM 20×CAG and 135 mM MgCl_2_; Lower panel: FRAP measurement of the droplet formed by 65 μM 20×CAG and 135 mM MgCl_2._ **d** Lowering the pH or adding EDTA dissipates the droplets. Scale bars are 10 μm in a and d, 5 μm in b and c.

To assess the fluidity of the RNA condensed phase, we conjugated a Cy5-dye at the 3’-end of the 20×CAG-repeat RNA, and doped the labeled RNA to the unlabeled RNA. The RNA droplets did not fuse after an extended incubation time (**Figure 1b**), while the Cy5 fluorescence shows little to small recovery after photobleaching (FRAP) at various locations of the selected droplets (**Figure 1c**). As such, though the RNA molecule used here has fewer CAG repeats than previously reported,^[5a]^ it also assembles to form largely immobilized droplets.

Though gel-like, the RNA droplets can be reversed and dissipated at an acidic pH (**Figure 1d**). On the other hand, increasing the concentration of Mg^2+^ can lead to the formation of larger RNA droplets (**Figure 1c** lower panel, and **Figure S2**). Conversely, the presence of a metal chelator, i.e., ethylenediaminetetraacetate (EDTA), nearly abolished the phase separation of the RNA molecules (**Figure 1d**). As such, the high-concentration of Mg^2+^ shields unfavorable electrostatic interactions between the RNA backbone, and consequently promotes intermolecular interactions.

When adding EDTA, we noticed that some of the droplets did not completely disappear, but instead morphed to a vesicle-like structure. Intrigued, we prepared the solutions with 86 μM RNA and with different concentrations of Mg^2+^. In the presence of 135 mM Mg^2+^, large droplets can be observed using differential interference contrast (DIC) microscopy (**Figure 2a**). In comparison, with lower concentrations of Mg^2+^ (60 mM and 35 mM) added to the same RNA solution, smaller RNA droplets are observed. Interestingly, in the presence of 35 mM Mg^2+^, the 20×CAG RNA molecules phase-separate to form hollow vesicle-like structure (**Figure 2a**, right panel).

**Figure 2.**
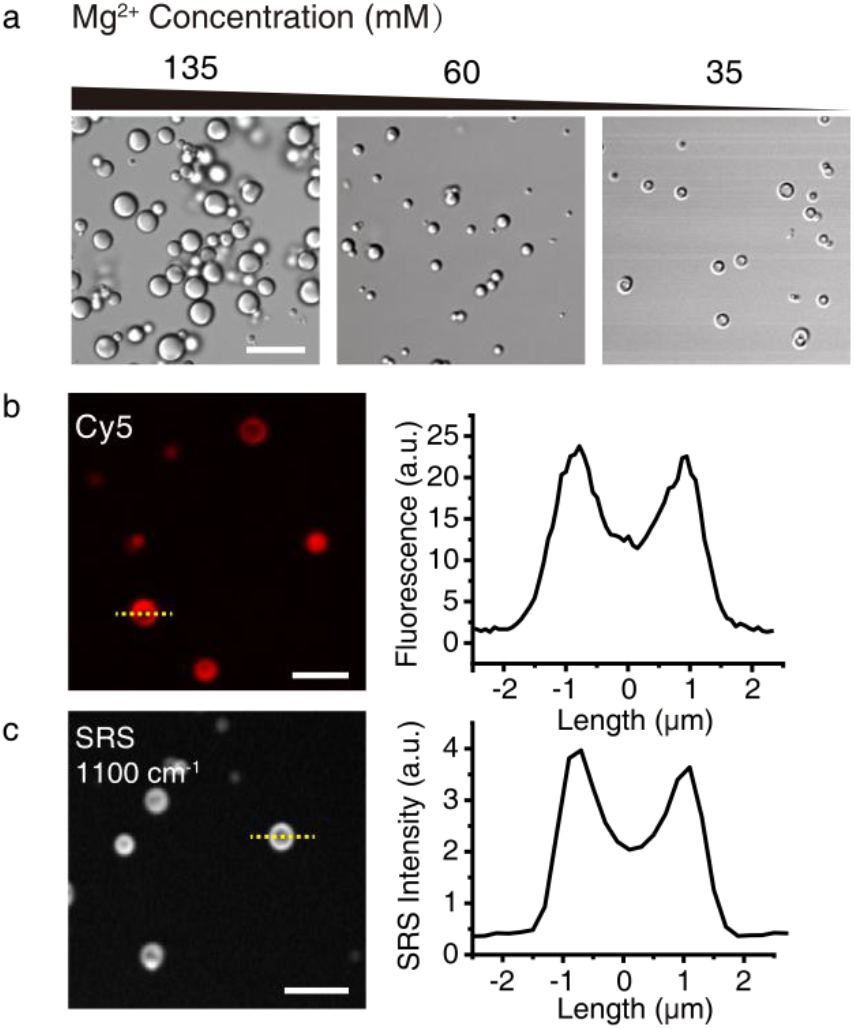
Formation of vesicle-like RNA droplets. **a** DIC images of droplets formed by 86 μM 20×CAG RNA at three Mg^2+^ concentrations. b Fluorescence intensity analysis of a hollow RNA droplet structure formed by 86 μM 20×CAG and 35 mM MgCl_2_, with Cy5-conjugated RNA was doped with unlabeled RNA. **c** Analysis of Raman spectral intensity at 1100 cm^−1^ of a selected droplet. The intensities were assessed along the dashed line, with a pixel resolution of 0.2 μm. Scale bars are 10 μm in a, 5 μm in b and c.

For those apparently hollow RNA droplets, fluorescence confocal microscopy imaging indicates that the RNA concentration is indeed lower at the center than at the rim (**Figure 2b**). To rule out aggregation-caused fluorescence quenching, we resorted to Raman spectroscopy. The Raman band at 1100 cm^−1^ arises from symmetric stretching of RNA phosphate backbone, and its intensity is proportional to the overall concentration of nucleobases and is insensitive to RNA structural changes.^[12a,17]^ The 1100 cm^−1^ Raman band intensity indicates that the RNA molecule is about twice as concentrated at the rim of the vesicles (**Figure 2c**), consistent with the fluorescence microscopy observation. The hollow RNA droplets remain gel-like, as multiple droplets can form a complex and multi-chamber structure (Supporting Information, **Figure S3**).

While keeping the Mg^2+^ concentration constant at 135 mM, we prepared the solutions of 20×CAG RNA at 35 μM and 65 μM, respectively. The RNA molecules can readily phase separate at both concentrations (**Figure 3a**, left and middle panels). Furthermore, at a higher RNA concentration (86 μM) but a lower Mg^2+^ concentration (35 mM), the RNA droplets are smaller and hollow (Figure 3a, right panel). As such, the morphology of the RNA condensate depends on the relative RNA/Mg^2+^ ratio. We name these three morphologies as type I, II, and III droplets. Assessed with the Raman intensity at 1100 cm^−1^, type-II RNA droplets are larger than type-I droplets, and have a higher enrichment of the RNA molecules (**Figure 3b,c**). The three types of RNA droplets have the diameters of 3.0±0.7 μm, 4.4±0.7 μm, and 2.4±0.7 μm, respectively (**Figure 3d**). Interestingly, though prepared from the highest RNA concentration, the type-III vesicle-like droplets are the smallest and have the lowest enrichment ratio (**Figure 3c**), which further attests the importance of Mg^2+^ in RNA coacervation.

**Figure 3.**
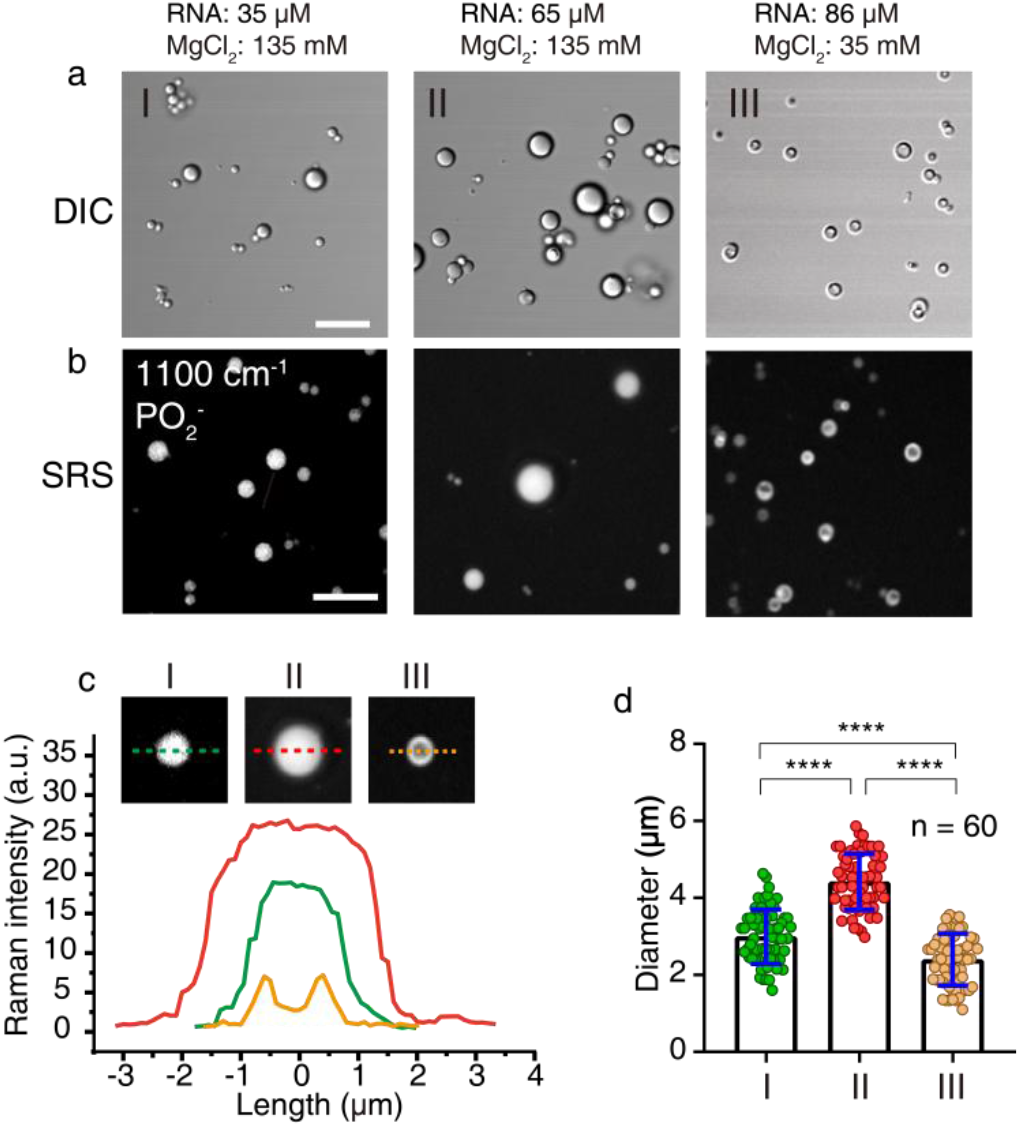
Morphology of the RNA droplets at different RNA/Mg^2+^ ratios. **a,b** DIC and SRM images at 1100 cm^−1^ for the RNA droplets formed with 35 μM RNA and 135 mM MgCl_2_ (morphology I), 65 μM RNA and 135 mM MgCl_2_ (morphology II), and 86 μM RNA and 35mM MgCl_2_ (morphology III). **c** The Raman intensities at 1100 cm^−1^ along the dashed lines for type-I, II and III droplets. **d** Statistics of the diameters of the RNA droplets for the three morphologies in DIC images. *n* = 60 and *p* < 0.0001. Scale bars, 10 μm.

To understand the molecular basis that leads to the formation of RNA condensate, we collected spontaneous Raman spectra of the 20×CAG RNA in the dilute and condensed phases. With the RNA concentration in the two phases normalized by the intensity of the Raman peak at 1100 cm^−1^, we assessed the relative intensities for other Raman bands. It has been established that upon the formation of an RNA duplex or hairpin, hypochromicity is observed for Raman spectral bands between 600 and 800 cm^−1^. In particular, hypochromicity for 727 cm^−1^ and 785 cm^−1^ bands have been attributed to the decrease of ring breathing in the base-paired RNA for adenine and cytosine nucleobases, respectively.^[12a,18]^ On the other hand, hyperchromicity is observed for Raman spectral bands between 1200 and 1600 cm^−1^. In particular, the 1250 cm^−1^, 1481 cm^−1^, and 1577 cm^−1^ bands^[17c,19]^ increase in intensities upon the formation of canonical RNA secondary structures, which is resulted from the increased ring stretching and collective vibration,^[12a,17c,20]^ as well as increased base stacking and clustering.^[12a,21]^

We show that, upon the coacervation of the repeat-expansion RNA molecules, the Raman intensities for 727 and 785 cm^−1^ bands are higher in the droplets than in the dilute phase (**Figure 4**). The 20×CAG RNA is predicted to form a stable hairpin structure, and may also hybridize to form a duplex (**Figure S4**) depending on the annealing condition. The increased Raman intensities at 727 cm^−1^ and 785 cm^−1^ indicate that the RNA in the condensed phase contains less base-paired structure. Specifically, the Raman intensities at 727 cm^−1^ and 785 cm^−1^ are 3.5-fold and 2.6-fold higher in the droplets, respectively, suggesting non-canonical base pairs involving adenines^[22]^ are more easily disrupted upon RNA coacervation. The increase of more single-stranded RNA regions may also be due to steric hindrance, which precludes proper base paring and the formation of RNA duplex or hairpin in a crowded environment.

**Figure 4.**
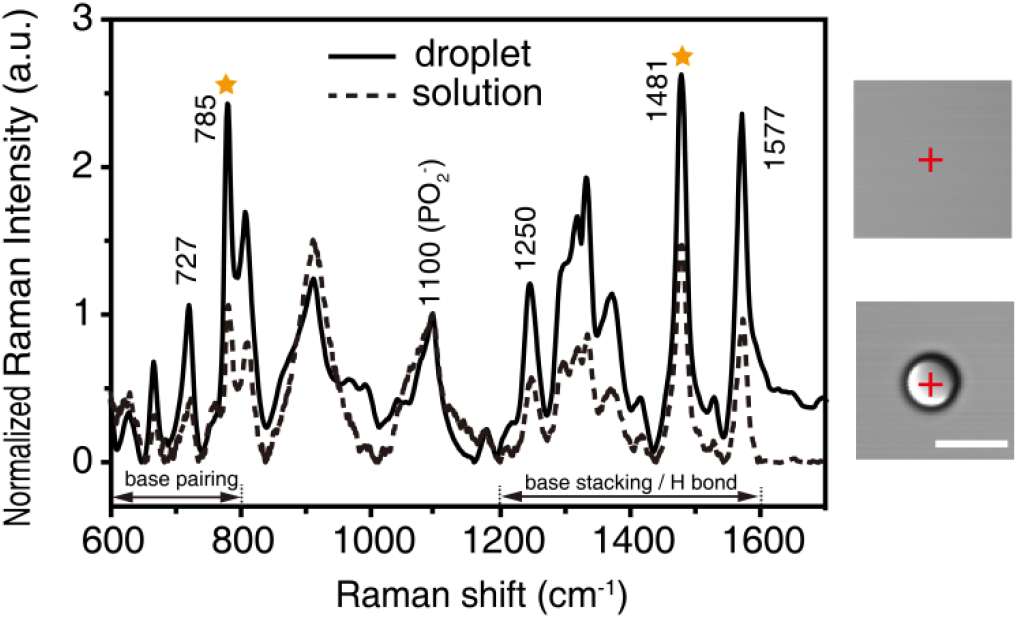
Spontaneous Raman Spectrum of a representative RNA droplet. The droplets were prepared with 65 μM 20×CAG repeat RNA,135 mM MgCl2 in the 10 mM Tris•HCl pH 7 buffer. As a control, the Raman spectrum was collected for the RNA solution at 1.7 mM with no annealing treatment. Scale bar, 5 μm. The starred peaks are further subjected to stimulated Raman spectroscopy analysis.

On the other hand, the Raman intensities at 1250 cm^−1^, 1481 cm^−1^, and 1577 cm^−1^ are higher in the condensed phase than in the dilute phase (**Figure 4**). As such, the nucleobases likely stack and cluster together in the droplets as opposed to the canonical hydrogen-bonding and base-pairing associated with the A-form duplex or hairpin structure. The stacking/clustering should involve stacking between RNA single-stranded regions^[23]^ or stacking between multiple nucleobases.^[24]^

To map the location-specific RNA structure in the condensed phase, we employed SRM^[13a]^ and collected a hyperspectral stack of Raman images at wavelengths from 650 to 1600 cm^−1^ for the three types of RNA droplets. After normalizing by the Raman intensity at 1100cm^−1^, the intensity differences at other wavelengths become obvious **(Figure S5)**. The Raman intensities at 785 cm^−1^ and 1481 cm^−1^ are just slightly higher at the center than at the rim of the type-I droplet (**Figure 5a,g**). In contrast, the Raman intensities at 785 cm^−1^ and 1481 cm^−1^ are lower at the center than at the rim of the type-II and type-III droplets. This means, at the center of these two types of RNA droplets, the RNA structure is more based-paired but less clustered (**Figure 5b,c,h,i**), akin the RNAs in a dilute phase.

**Figure 5.**
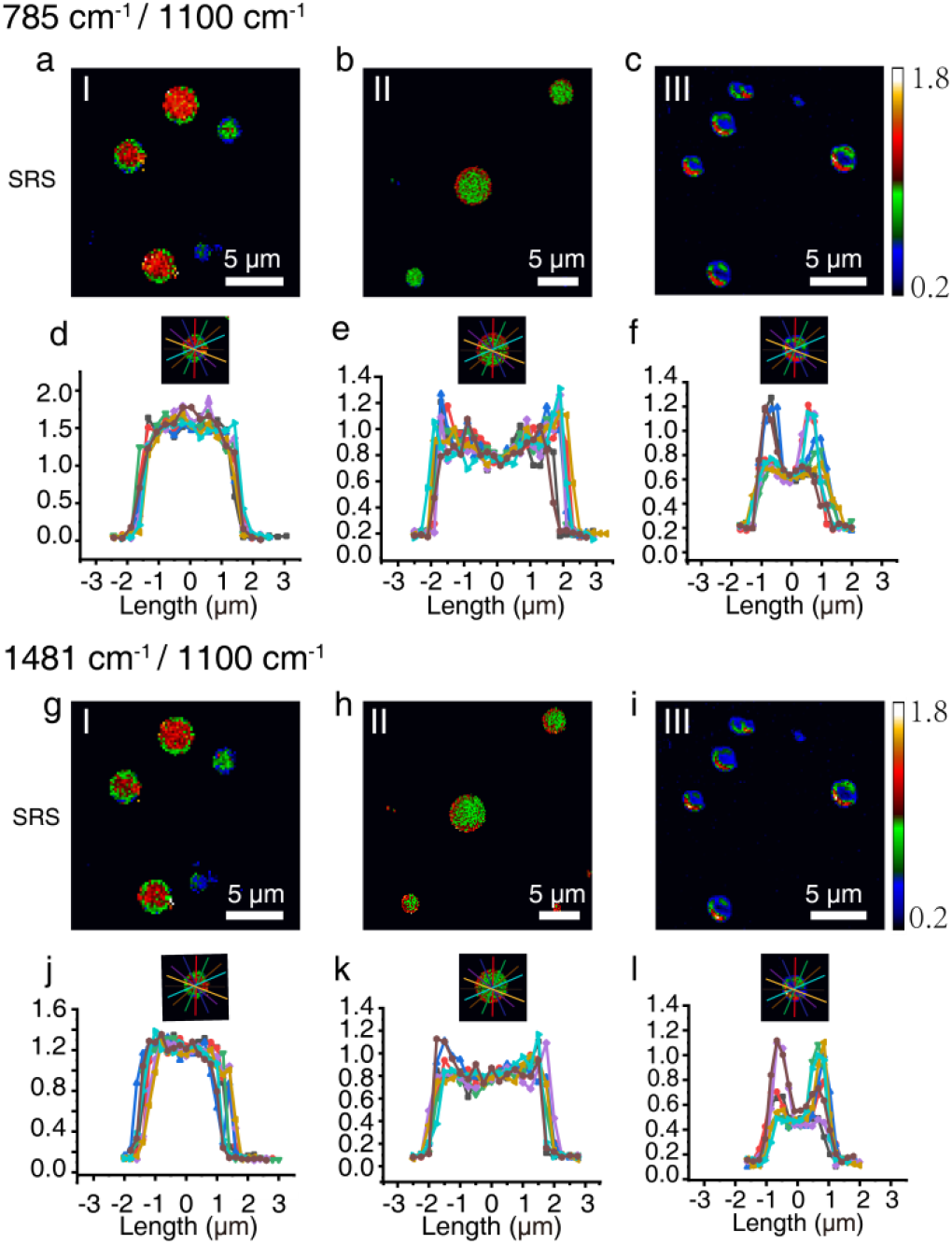
Hyperspectral stimulated Raman scattering microscopy analysis of the three types of RNA droplets. **a-c, g-i** SRM images at 785 cm^−1^ and 1480 cm^−1^, respectively. **d-f, j-l** Quantitative assessment of the inhomogeneous distribution of the RNA structures within the droplets. Each line indicates an arbitrary line across a selected droplet at 785 cm^−1^ and 1480 cm^−1^, respectively. The intensities are normalized to Raman intensity at 1100 cm^−1^. Scale bars, 5 μm.

Further, we measured the radial distributions of Raman intensities at various wavelengths (**Figure S6**). For the type-I droplets, the 758 cm^−1^ and 1481 cm^−1^ Raman intensities at the rim are nearly comparable to those at the center, meaning a quite homogenous architecture (**Figure 5d, j**). On the other hand, both type-II and type-III droplets have well-defined rims, which are characterized by sharp transitions in Raman intensities (**Figure 5e,f,k,l**). As such, the type-II droplets also form a vesicle-like structure, even though they do not appear so under the light microscopy. Thus, the hollowing of RNA droplets correlates with the increase of RNA/Mg^2+^ ratio, and the RNA molecules tend to cluster and stack at the rims of both type-II and type-III droplets (**Figure 6**).

**Figure 6.**
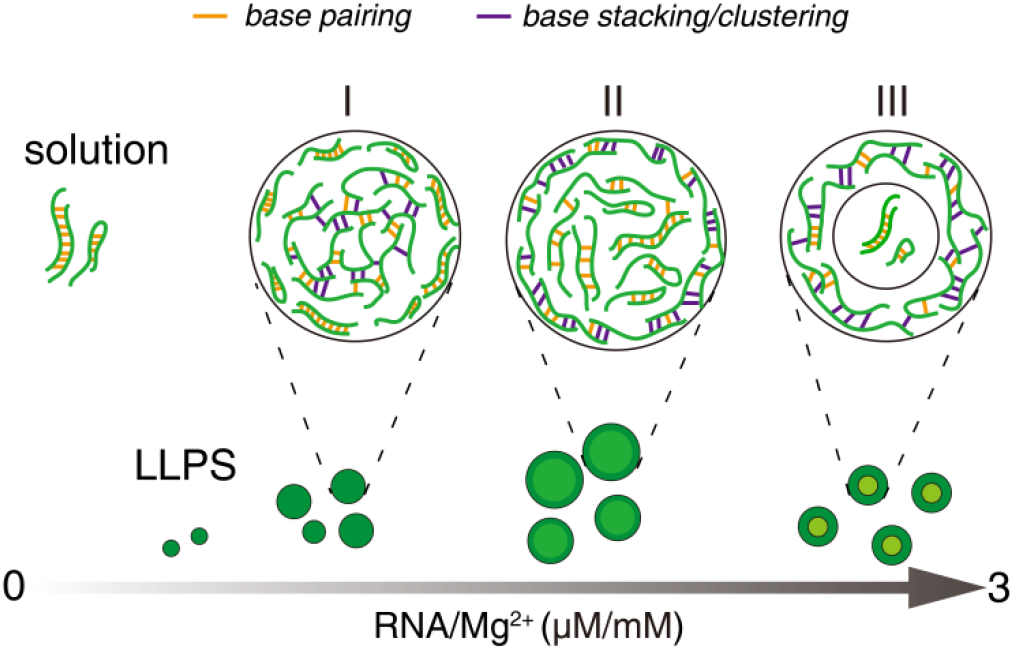
Schematic for the different structures of RNA droplets formed by 20×CAG at different RNA/Mg^2+^ ratios. When the RNA concentration increases or Mg^2+^ concentration decreases, the RNA molecules are more likely to phase-separate into type-II and type-III droplets, with the outer rim containing a large proportion of stacked/clustered nucleobases.

In summary, we have used SRM to uncover the structural features of a condensed phase, homotypically formed by a 20×CAG repeat RNA molecule. The addition of a large excess of divalent cation neutralizes electrostatic repulsions between RNA molecules, as previously shown.^[16]^ Significantly, Raman spectroscopic analysis indicates that the stacking and clustering of the nucleobases, instead of hydrogen-bonding and canonical base pairing, contribute to the formation of the RNA droplets (**Figure 4**). Indeed, partial protonation of the N^3^ atom of cytosine and N^1^ atom of adenine nucleobases at an acidic pH^[25]^ would disrupt nucleobase clustering and dissipates RNA droplets, as we show in **Figure 1d**.

Depending on the RNA/Mg^2+^ ratio, we also show that the RNA droplets adopt different architectures and morphologies. When the Mg^2+^ concentration decreases, the repulsion between the phosphate backbone increases. With less counterion present, the RNA droplets become gradually hollowed, while nucleobase stacking and clustering, a form of hydrophobic interactions, become more dominant for the formation of RNA droplets. Ultimately, the RNA molecules form a vesicle-like structure, with the center containing much fewer molecules and a less nucleobase-clustered structure (**Figure 6**). In comparison to base pairing, nucleobase clustering can be more restrictive, affording a large network of intermolecular interactions. Indeed, the FRAP experiment reveals a partial recovery of the fluorescence for the type-II droplets (**Figure 1c**, lower panel, and **Figure S7**), corroborating the SRM results.

Our finding thus has broader implications in the active research field of phase separation. When co-phase separated with proteins, RNA molecules are often found absorbed at the surface of those droplets.^[4d,26]^ Moreover, the RNA component has been shown to limit the size of RNA-protein condensate.^[27]^ Since the interwoven nucleobase-clustered RNA assembly is likely mechanically rigid, a protective RNA shell would define the physical boundaries of membrane-less organelles. In this regard, the repeat-expansion RNA molecules act as a Pickering agent,^[28]^ as recently ascribed for MEG-3 protein absorbed at the surface of P granules.^[29]^

## Acknowledgements

The work was supported by the National Key R&D Program of China (2018YFA0507700 to C.T.) and the National Natural Science Foundation of China (31971155 to Z.G.). P.W. acknowledges the supports from the National Natural Science Foundation of China 62075076 and Innovation Fund of the Wuhan National Laboratory for Optoelectronics.

## Entry for the Table of Contents

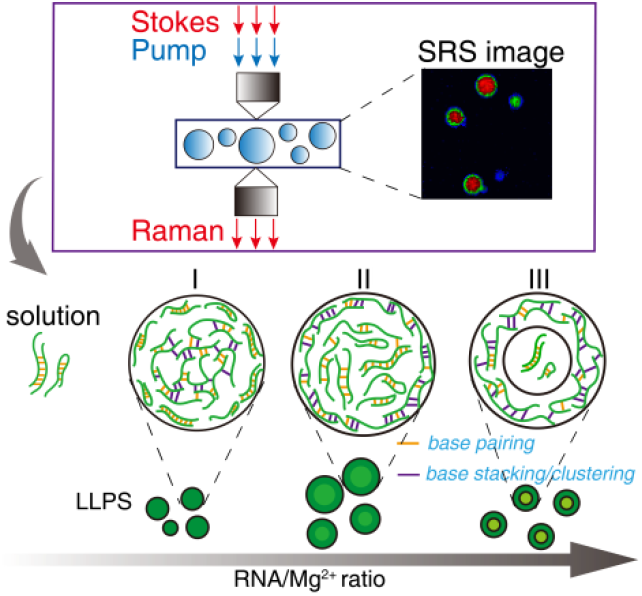

Phase separation of repeat expansion RNA molecules leads to the formation of gel-like droplets. Ma et. al showed that depending on the RNA/Mg^2+^ ratio, a 20×CAG repeat RNA forms small, large, and hollow droplets. Using hyperspectral stimulated Raman spectroscopy, they demonstrated that nucleobase clustering is a driving force for RNA coacervation, and is predominant at the rim of large and hollow droplets.

Institute Twitter username: @PKU_1898

Researcher Twitter username: @Chun Tang

## References

[1] (a) P. Anderson, N. Kedersha, Nat. Rev. Mol. Cell Biol. 2009, 10, 430–436; (b) J. Berry, S. C. Weber, N. Vaidya, M. Haataja, C. P. Brangwynne, Proc Natl Acad Sci U S A 2015, 112, 5237–5245; (c) B. Tsang, J. Arsenault, R. M. Vernon, H. Lin, N. Sonenberg, L. Y. Wang, A. Bah, J. D. Forman-Kay, Proc Natl Acad Sci U S A 2019, 116, 4218–4227; (d) T. J. Nott, E. Petsalaki, P. Farber, D. Jervis, E. Fussner, A. Plochowietz, T. D. Craggs, D. P. Bazett-Jones, T. Pawson, J. D. Forman-Kay, A. J. Baldwin, Mol. Cell 2015, 57, 936–947; (e) H. Zhang, X. Ji, P. L. Li, C. Liu, J. Z. Lou, Z. Wang, W. Y. Wen, Y. Xiao, M. J. Zhang, X. L. Zhu, Sci China Life Sci 2020, 63, 953–985; (f) M. M. Fay, P. J. Anderson, J. Mol. Biol. 2018, 430, 4685–4701; (g) M. Polymenidou, Science 2018, 360, 859–860.

[2] (a) A. Molliex, J. Temirov, J. Lee, M. Coughlin, A. P. Kanagaraj, H. J. Kim T. Mittag, J. P. Taylor, Cell 2015, 163, 123–133; (b) Y. Shin, C. P. Brangwynne, Science 2017, 357.

[3] M. Fuxreiter, M. Vendruscolo, Nat. Cell Biol. 2021, 23, 587–594.

[4] (a) P. R. Banerjee, A. N. Milin, M. M. Moosa, P. L. Onuchic, A. A. Deniz, Angew. Chem. Int. Ed. 2017, 56, 11354–11359; (b) S. Maharana, J. Wang, D. K. Papadopoulos, D. Richter, A. Pozniakovsky, I. Poser, M. Bickle, S. Rizk, J. Guillen-Boixet, T. M. Franzmann, M. Jahnel, L. Marrone, Y. T. Chang, J. Sterneckert, P. Tomancak, A. A. Hyman, S. Alberti, Science 2018, 360, 918–921; (c) S. Boeynaems, A. S. Holehouse, V. Weinhardt, D. Kovacs, J. Van Lindt, C. Larabell, L. Van Den Bosch, R. Das, P. S. Tompa, R. V. Pappu, A. D. Gitler, Proc Natl Acad Sci U S A 2019, 116, 7889–7898; (d) I. Alshareedah, M. M. Moosa, M. Raju, D. A. Potoyan, P. R. Banerjee, Proc Natl Acad Sci U S A 2020, 117, 15650–15658.

[5] (a) A. Jain, R. D. Vale, Nature 2017, 546, 243–247; (b) M. M. Fay, P. J. Anderson, P. Ivanov, Cell Rep. 2017, 21, 3573–3584; (c) Y. Teng, H. Tateishi-Karimata, N. Sugimoto, Biochemistry 2020, 59, 1972–1980.

[6] P. C. Bevilacqua, A. M. Williams, H. L. Chou, S. Assmann, RNA 2021, pp 1–28.

[7] S. Peng, W. Li, Y. Yao, W. Xing, P. Li, C. Chen, Proc Natl Acad Sci USA 2020, 117, 27124–27131.

[8] (a) S. Elbaum-Garfinkle, Y. Kim, K. Szczepaniak, C. C. H. Chen, C. R. Eckmann, S. Myong, C. P. Brangwynne, Proc Natl Acad Sci USA 2015, 112, 7189–7194; (b) I. Nasir, P. L. Onuchic, S. R. Labra, A. A. Deniz, BBA Proteins and Proteomics 2019, 1867, 980–987.

[9] (a) J. P. Brady, P. J. Farber, A. Sekhar, Y. H. Lin, R. Huang, A. Bah, T. J. Nott, H. S. Chan, A. J. Baldwin, J. D. Forman-Kay, L. E. Kay, Proc Natl Acad Sci U S A 2017, 114, 8194–8203; (b) A. C. Murthy, G. L. Dignon, Y. Kan, G. H. Zerze, S. H. Parekh, J. Mittal, N. L. Fawzi, Nat. Struct. Mol. Biol. 2019, 26, 637–648.

[10] K. Murakami, S. Kajimoto, D. Shibata, K. Kuroi, F. Fujii, T. Nakabayashi, Chem. Sci. 2021, 12, 7411–7418.

[11] Y. Li, S. Liu, H. Ni, H. Zhang, H. Zhang, C. Chuah, C. Ma, K. S. Wong, J. W. Y. Lam, R. T. K. Kwok, J. Qian, X. Lu, B. Z. Tang, Angew. Chem. Int. Ed. 2020, 59, 12822–12826.

[12] (a) A. J. Hobro, D. M. Standley, S. Ahmad, N. I. Smith, Phys. Chem. Chem. Phys. 2013, 15, 13199–13208; (b) A. L. Wilson, C. Outeiral, S. E. Dowd, A. J. Doig, P. L. A. Popelier, J. P. Waltho, A. Almond, Chemi. Commun. 2020, 3.

[13] (a) M. C. Wang, W. Min, C. W. Freudiger, G. Ruvkun, X. S. Xie, Nature Methods 2011, 8, 135–152; (b) Y. Bi, C. Yang, Y. Chen, S. Yan, G. Yang, Y. Wu, G. Zhang, P. Wang, Light Sci. Appl. 2018, 7.

[14] (a) M. Ji, M. Arbel, L. Zhang, C. W. Freudiger, S. S. Hou, D. Lin, X. Yang, B. J. Bacskai, X. S. Xie, Sci. Adv. 2018, 4; (b) D. Zhang, P. Wang, M. N. Slipchenko, J. X. Cheng, Acc. Chem. Res. 2014, 47, 2282–2290.

[15] L. Wei, F. Hu, Z. Chen, Y. Shen, L. Zhang, W. Min, Acc. Chem. Res. 2016, 49, 1494–1502.

[16] P. L. Onuchic, A. N. Minn, I. Alshareedah, A. A. Deniz, P. R. Banerjee, Sci. Rep. 2019, 9.

[17] (a) B. Gong, J. H. Chen, R. Yajima, Y. Y. Chen, E. Chase, D. M. Chadalavada, B. L. Golden, P. R. Carey, P. C. Bevilacqua, Methods 2009, 49, 101–111; (b) P. Carmona, M. Molina, A. Rodriguez-Casado, J Raman Spectrosc 2009, 40, 893–897; (c) J. Morla-Folch, H. N. Xie, R. A. Alvarez-Puebla, L. Guerrini, Acs Nano 2016, 10, 2834–2842.

[18] M. C. Chen, G. J. Thomas, Biopolymers 1974, 13, 615–626.

[19] Y. Li, T. Gao, G. Xu, X. Xiang, B. Zhao, X. X. Han, X. Guo, Anal. Chem. 2019, 91, 7980–7984.

[20] (a) G. J. Thomas, M. C. Chen, K. A. Hartman, Biochimica et Biophysica Acta 1973, 324, 37–49; (b) A. J. Hobro, M. Rouhi, E. W. Blanch, G. L. Conn, Nucleic Acids Res. 2007, 35, 1169–1177.

[21] B. Hernandez, V. Baumruk, N. Leulliot, C. Gouyette, T. Huynh-Dinh, M. Ghomi, J. Mol. Struct. 2003, 651, 67–74.

[22] N. B. Leontis, J. Stombaugh, E. Westhof, Nucleic Acids Res. 2002, 30, 3497–3531.

[23] B. Van Treeck, R. Parker, Cell 2018, 174, 791–802.

[24] A. M. Williams, R. R. Poudyal, P. C. Bevilacqua, Biochemistry 2021, 60, 2715–2726.

[25] (a) R. Krishnamurthy, Acc. Chem. Res. 2012,45, 2035–2044; (b) P. Thaplyal, P. C. Bevilacqua, Chapter Nine - Experimental Approaches for Measuring pKa’s in RNA and DNA. In Methods in Enzymology, Burke-Aguero, D. H., Ed. Academic Press, 2014, pp 189–219.

[26] A. Cochard, G. J. Navarro, S. Kashida, M. Kress, Z. Gueroui, 2021.

[27] M. Garcia-Jove Navarro, S. Kashida, R. Chouaib, S. Souquere, G. Pierron, D. Weil, Z. Gueroui, Nat. Commun. 2019, 10, 3230.

[28] A. B. Pawar, M. Caggioni, R. W. Hartel, P. T. Spicer, Faraday Discuss. 2012, 158, 341–350.

[29] A. W. Folkmann, A. Putnam, C. F. Lee, G. Seydoux, Science 2021, 373, 1218–1224.

