## Supplemental Figure S4 for "Nucleobase clustering contributes to the formation and hollowing of repeat-expansion RNA condensate"

©Wiley-VCH 2021

69451 Weinheim, Germany

**Abstract:** RNA molecules with repeat expansion sequences can phase separate into gel-like condensate, and this process may lead to neurodegenerative diseases. Here we report that in the presence of Mg^2+^ ion, RNA molecules containing 20×CAG repeats coacervate into filled droplets or hollowed condensate. Using hyperspectral stimulated Raman spectroscopy, we show that RNA coacervation is accompanied by the stacking and clustering of nucleobases, while forfeiting the canonical base-paired structure. At an increasing RNA/Mg^2+^ ratio, the RNA droplets first expand in sizes, and then shrink and adopt hollow vesicle-like structures. Significantly, for both large and vesicle-like droplets, the nucleobase-clustered structure is more prominent at the rim than at the center, accounting for the rigidification of RNA droplets. Thus, our finding has broad implications for the general aging processes of RNA-containing membrane-less organelles.

DOI: 10.1002/anie.2021XXXXX

Experimental Procedures

**RNA sample preparation.** The RNA molecule with 20×CAG repeat sequence was prepared with solid phase synthesis on an ABI-3400 oligonucleotide synthesizer. Phosphoramidites of adenosine, guanosine, and cytosine were purchased from GenePharma, Shanghai, China. The separation of synthetic RNA from controlled pore glass (CPG) and the removal of various protecting groups followed an established protocol.^[1]^ Subsequently, the RNA sample was precipitated with pre-chilled butyl alcohol, purified on a Source Q column (GE Healthcare), and concentrated and buffer-exchanged with Amicon Ultra (5K MWCO, Millipore). For the synthesis of labeled 20×CAG, 3'-Cy5 CPG (GenePharma) were used. The purity of RNA was evaluated by electrophoresis on a 15% polyacrylamide denaturing gel.

**Confocal fluorescence microscopy imaging.** The RNA molecule with 20×CAG repeat sequence was prepared in the 10 mM pH 7.0 Tris•HCl buffer containing 25 mM NaCl buffer. Cy5-conjugated RNA and unlabeled RNA were mixed at 1:25 ratio. The RNA molecules were heated to 95℃ for 3 min in a metal bath, and slowly cooled down to 25 ºC. RNA samples were placed in a small parafilmed chamber molded between two cover glasses. Formation of the RNA droplets were visualized at room temperature using an A1 confocal laser scanning microscope (Nikon), equipped with a 60×1.42 numerical aperture oil objective. The differential interference contrast (DIC) images were obtained with the excitation of a 561 nm laser, and the fluorescence images were obtained at 640 nm. For fluorescence recovery after photobleaching (FRAP) experiments, a region of ~0.8 μm^2^ within a droplet formed by the fluorophore-doped RNA was photobleached using a 561 nm laser with 100% laser power. The droplet was then imaged continuously to monitor the recovery process over time.

**Spontaneous Raman spectroscopy.** The Raman spectra of the RNA molecules were recorded with 532 nm excitation (LabRAM HR800, Horiba Jobin Yvon). The Raman spectra of the dilute phase was obtained for an RNA solution at 1.7 mM concentration without annealing. The Raman spectra were measured from 500 to 2000 cm^-1^, and were de-baselined using the software LabSpec 5 (HORIBA Scientific).

**Hyperspectral stimulated Raman scattering microscopy.** The hyperspectral SRS imaging system is home-built with the use of a dual-output femtosecond laser (InSight DeepSee, Spectra-Physics, America). The detailed setup of the equipment has been reported previously,^[^^2^^]^ as shown in Figure S1. The laser has a synchronized pump (680-1300 nm, ~120 fs) and Stokes (1040 nm, ~220 fs) laser beams at a repetition rate of 80 MHz. The Stokes beam was modulated by a resonant electro-optical modulator (EO-AM-R-C2, Thorlabs) at 10.55 MHz for shot-noise limited detection. The pump beam was spatially overlapped with Stokes beam by a dichroic mirror (DMSP1000L, Thorlabs). Pump and Stokes pulses were chirped to ~3 ps by a 64 cm long SF57 glass rod. By turning the relative delay time, Raman wave-number was spectrally focused for the hyperspectral SRS imaging. The two laser beams were fed to a laser scanning microscope equipped with a two-axis galvanometer (GVS002, Thorlabs). A 60× water immersion objective (Olympus) was used for imaging. The transmitted pump beam was collected with a high numerical aperture oil condenser, and detected with a photodiode (S3994-01, Hamamatsu). Two high-optical-density short-pass filters (ET980SP, Chroma, America) were used to block the Stokes beam. The output current of the photodiode was amplified by a home-built 10.55 MHz resonant amplifier, and was demodulated by a digital lock-in amplifier (LIA, HF2LI, Zurich Instrument).

All laser powers were measured before objective (70% transmission), and the dwell time was set at 10 µs. For all hyperspectral SRM imaging, the laser power of pump and Stokes beams was set to 60 and 160 mW, respectively. The SRS spectrum was acquired by adjusting the delay time of pump laser and Stokes laser. For the SRM image of droplets formed by 35 µM 20×CAG 135 mM MgCl_2_, the field of view (FOV) is 100×100 μm^2^ with 300×300 pixels. For the SRM image of the droplets formed by 65 µM 20×CAG 135 mM MgCl_2_, the FOV is 60×60 μm^2^ with 300×300 pixels. For the SRM image of shell formed by 86 µM 20×CAG 35 mM MgCl_2_, the FOV is 60×60 μm^2^ with 300×300 pixels. A hyperspectral stack of 320 images ranging from 650 to 1600 cm^−1^ was obtained by adjusting the laser wavelength to 964 nm, 934 nm, 916 nm and 900 nm, respectively. The spectral intensities were normalized by the intensity of 1100 cm^-1^ band.

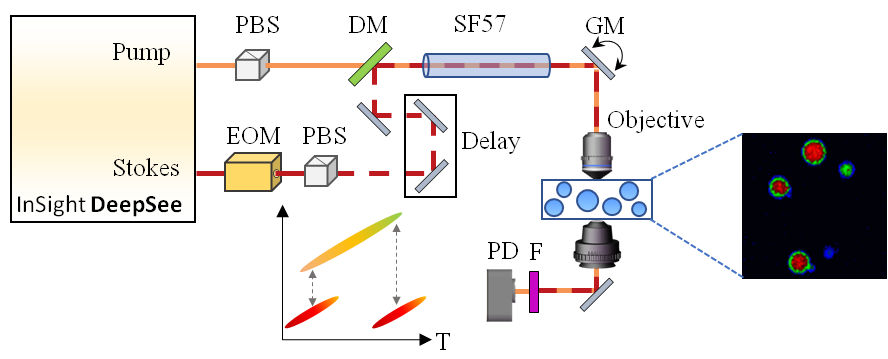

Figure S1. Schematic of the spectral-focusing SRM imaging system. PBS: Polarizing Beamsplitter; EOM: electro-optic modulator; DM: dichroic mirror; GM: galvanometer; F: optical filters; PD: photodiode

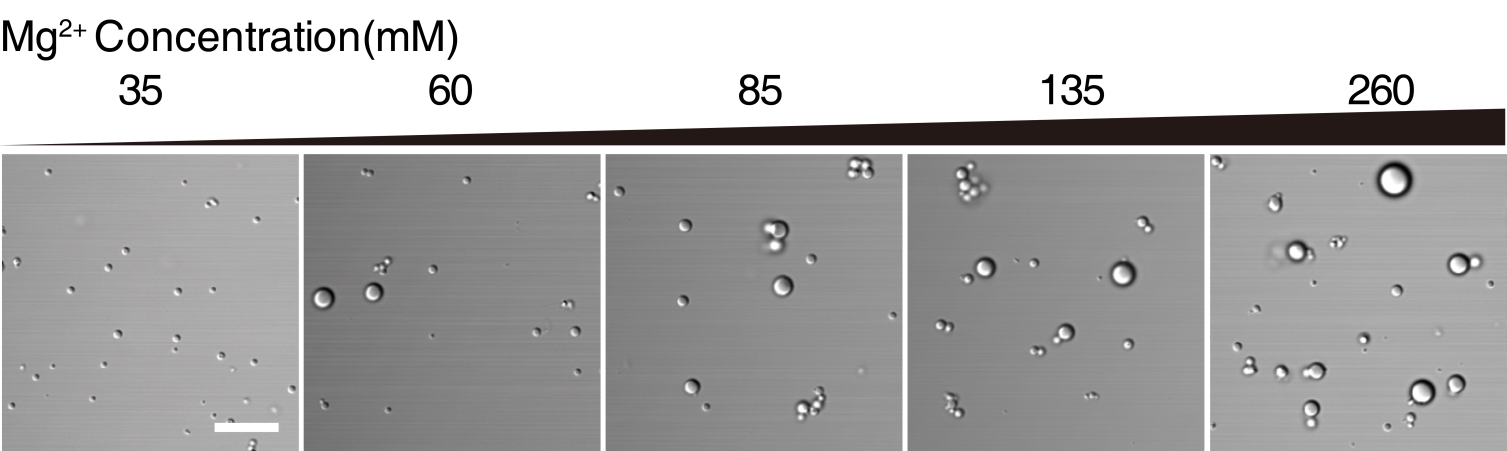

**Figure S2.** DIC images of RNA droplets formed at an increasing concentration of MgCl_2_. The initial sample contains 35 µM 20×CAG RNA in 10 mM pH 7 Tris•HCl buffer. Scale bars, 10 µM.

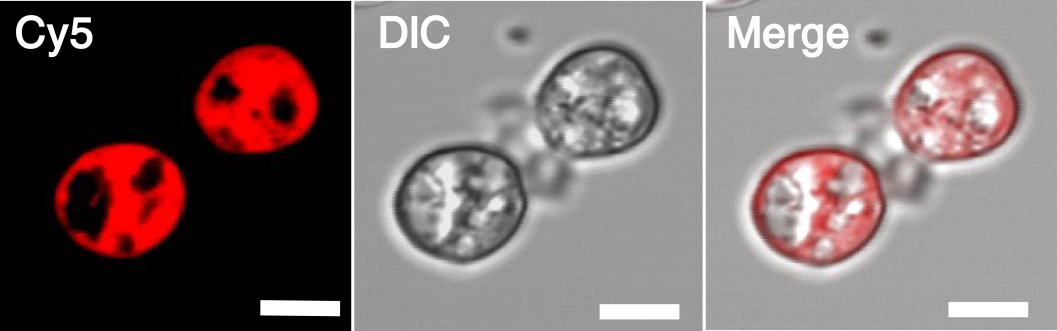

**Figure S3.** Partial fusion of the hollow RNA liquid droplets, with characteristic multi-chambered structure. The initial sample contains 86 µM 20×CAG RNA in the presence of 35 mM MgCl_2_. Scale bars, 5 µm.

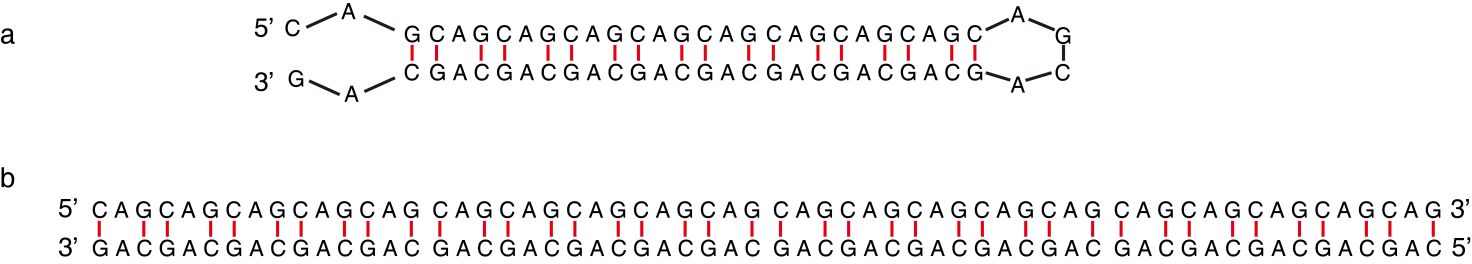

**Figure S4. a** The hairpin structure of 20×CAG RNA predicted by Mfold (http://www.unafold.org/mfold/applications/rna-folding-form.php), with predicted stabilization energy ΔG = -19.60 kcal/mol ; **b** The duplex structure formed by by 20×CAG RNA, with predicted stabilization energy ΔG = -41.3 kcal/mol.

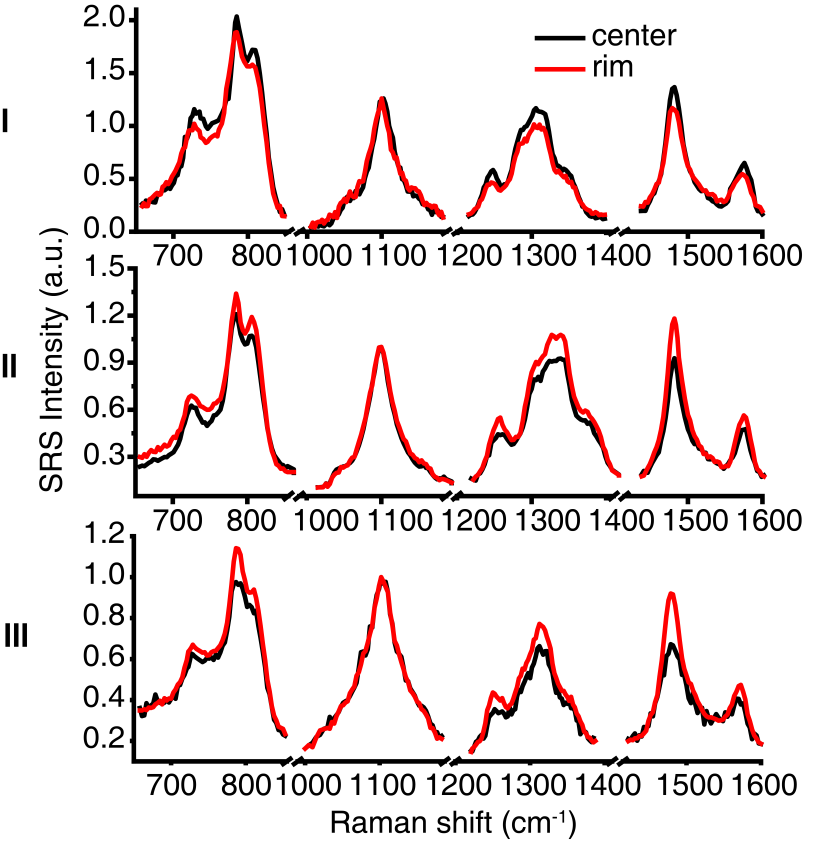

**Figure S5.** Normalized SRS spectrum of type-I, type-II and type-III droplets. The red line represents the spectrum at the rim of the droplets, the black represents the spectrum at the center of droplets.

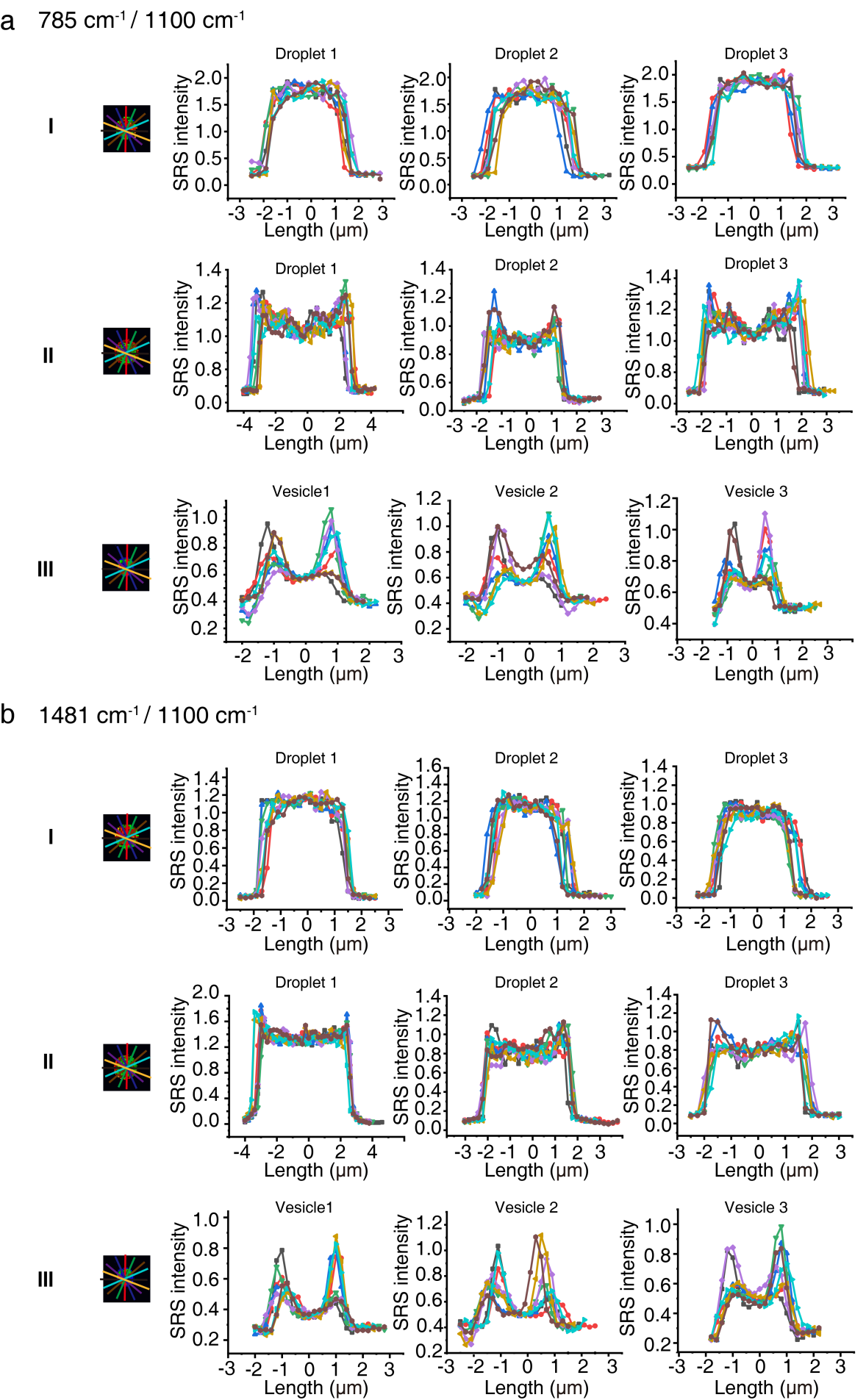

**Figure S6.** SRM analysis of the architecture of RNA droplets. **a** Quantitative assessment of the inhomogeneous distribution of the RNA structures within the type-I, type-II and type-III droplets at 785 cm^-1^. **b** Quantitative assessment of the inhomogeneous distribution of the RNA structures within the type-I, type-II and type-III droplets at 1480 cm^-1^. Each line indicates an arbitrary line across a selected droplet. The intensities are normalized to Raman intensity at 1100 cm^-1^.

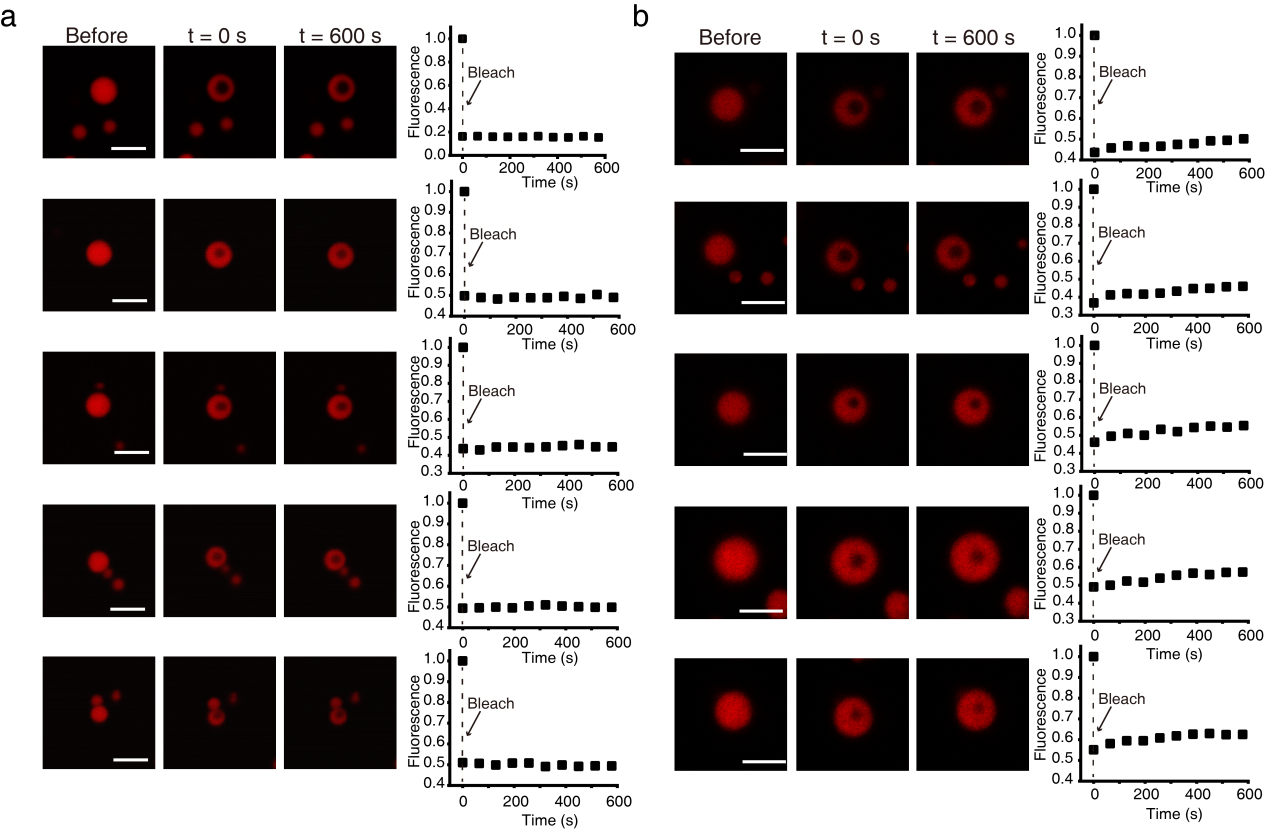

**Figure S7.** FRAP measurement of the type-I and type-II droplets. **a** FRAP measurement of the type-I droplet formed by 35 µM 20×CAG and 135 mM MgCl_2_. **b** FRAP measurement of the type-II droplet formed by 65 µM 20×CAG and 135 mM MgCl_2._ Scale bars, 5 µm.

### References

[1]	a) F. Wincott, A. DiRenzo, C. Shaffer, S. Grimm, D. Tracz, C. Workman, D. Sweedler, C. Gonzalez, S. Scaringe, N. Usman, *Nucleic Acids Res.* **1995**, 23, 2677-2684; b) Z. Gong, S. Yang, X. Dong, Q. F. Yang, Y. L. Zhu, Y. Xiao, C. Tang, *J. Mol. Biol.* **2020**, 432, 4523-4543.

[2]	a) S. Yan, S. Cui, K. Ke, B. Zhao, X. Liu, S. Yue, P. Wang, *Anal. Chem.* **2018**, 90, 6362-6366; b) S. Tian, H. Li, Z. Li, H. Tang, M. Yin, Y. Chen, S. Wang, Y. Gao, X. Yang, F. Meng, J. W. Lauher, P. Wang, L. Luo, *Nat. Commun.* **2020**, 11, 81.
